# Does impedance matter when recording spikes with polytrodes?

**DOI:** 10.1101/270058

**Authors:** Joana P. Neto, Pedro Baião, Gonçalo Lopes, João Frazão, Joana Nogueira, Elvira Fortunato, Pedro Barquinha, Adam R. Kampff

## Abstract

Extracellular microelectrodes have been widely used to measure brain activity, yet there are still basic questions about the requirements for a good extracellular microelectrode. One common source of confusion is how an electrode’s impedance affects the amplitude of extracellular spikes and background noise.

Here we discuss how an electrode’s impedance affects data quality in extracellular recordings, which is crucial for both the detection of spikes and their assignment to the correct neurons. This study employs commercial polytrodes containing 32 electrodes (177 μm^2^) arranged in a dense array. This allowed us to directly compare, side-by-side, the same extracellular signals measured by modified low impedance (~100 kOhm) microelectrodes with unmodified high impedance (~1 MOhm) microelectrodes. We begin with an evaluation of existing protocols to lower the impedance of the electrodes. The poly(3,4-ethylenedioxythiophene)-polystyrene sulfonate (PEDOT-PSS) electrodeposition protocol is a simple, stable, and reliable method for decreasing the impedance of a microelectrode up to tenfold. We next record *in vivo* using polytrodes that are modified in a ‘chess board’ pattern, such that the signal of one neuron is detected by multiple coated and non-coated electrodes. The performance of the coated and non-coated electrodes is then compared on measures of background noise and amplitude of the detected action potentials.

If the proper recording system is used, then the impedance of a microelectrode within the range of standard polytrodes (~ 0.1 to 2 MOhm) does not significantly affect data quality and spike sorting. This study should encourage neuroscientists to stop worrying about one more unknown.

## INTRODUCTION

Throughout the electrophysiology literature, an electrode’s impedance magnitude measured at 1 kHz in a saline solution is regularly used as a proxy for its ability to detect the activity of individual neurons [1]–[3]. The impedance is a measure of the ability of the electrode-solution interface circuit to resist the flow of charge across the interface’s phases (i.e., from the ionic to electronic conductor).

Do high impedance electrodes reduce the amplitude of the signal and/or increase the background noise? Clearly, lowering the signal-to-noise ratio (SNR) will make spike detection and sorting more difficult. How exactly does electrode impedance affect SNR? Several studies have shown an impact of electrode impedance on data quality [4]–[14]. However, there is also literature showing that electrode impedance does not affect the extracellular spikes recorded [15]–[17].

Commercially available silicon probes, also called polytrodes, have relatively high impedance electrodes due to their low surface area and small diameters (< 50 μm), which are suitable for recording single unit activity. Materials such as Au, Pt, and Ir are often used as the electrode material in polytrodes, and lowering the electrode impedance prior to recording is a ‘standard’ step in various laboratories [17]. How does one lower the impedance of commercial polytrodes?

Electrodeposition is a simple and reproducible technique, yet has great flexibility to produce a variety of coatings [4]. For more details about electrodeposition techniques see [18]. By electroplating Au or Pt, the surface roughness increases and the electrode impedance decreases [4], [17], [19], [20]. Over the last decade, conductive polymers, particularly poly(3,4-ethylenedioxythiophene) (PEDOT), have been electrodeposited onto electrodes due to their chemical stability and mechanical integrity when implanted in the brain [5], [6], [21]. Moreover, when compared to metals, these polymers are typically softer materials offering a more intimate contact between the electrode surface and brain tissue [22]. Prior to the electrodeposition, a dopant is added to the synthesis solution to improve conductivity; the most common dopant molecule is polystyrene sulfonate (PSS) [23], [24].

Our goal was simply to answer the question: ‘should I reduce the impedance of my polytrode electrodes’? Despite the prevalence of this question in the field, a definitive answer is still lacking. It is important to understand the impact of a particular electrode impedance and electrodeposition technique to determine if the effort to reduce impedance is necessary.

## EXPERIMENTAL SECTION

### Polytrodes

All experiments were performed with a commercially available 32-channel probe (A1×32-Poly3-5mm-25s-177-CM32, NeuroNexus), with 177 μm^2^ area electrodes (iridium) and an inter-site pitch of 22-25 μm (Figure S1 from Supplementary Information). Following each surgery, cleaning was performed by immersing the probe in a trypsin solution (Trypsin-EDTA (0.25 %), phenol red, TermoFisher Scientific) for 30-120 minutes and rinsing with distilled water [25].

### Coatings

NanoZ hardware and software (Neuralynx) was used to perform gold and PEDOT-PSS coating depositions. Moreover, both coatings were galvanostatically deposited in a two electrode cell configuration consisting of the probe microelectrodes individually selected as the working electrode and a platinum wire as the reference electrode. The reference wire was placed around the deposition cup while the probe was maintained at a fixed and equal distance to all points of the reference wire. By selecting ‘Manual Control’ from the NanoZ software it is possible to select individual probe electrodes [26].

For the gold coatings, a commercial non-cyanide gold solution was obtained from Neuralynx. The deposition solution for PEDOT-PSS consisted of 0.01 M of EDOT (Sigma-Aldrich, 97 %, Mw = 142.18) and 0.1 M of PSS (Sigma-Aldrich, Mw = 1000000) dissolved in deionized water. The optimal deposition parameters were −30 nA during 120 seconds for gold and +30 nA during 5 seconds for PEDOT-PSS [26]. Before and after the deposition, an electrode’s impedance magnitude at 1 kHz, in sterile phosphate buffer saline solution (PBS, 1 mM, pH 7.4), was measured with the NanoZ. Post-deposition assessment of coating morphology was performed by scanning electron microscopy (SEM-FIB, Zeiss Auriga).

### Electrochemical characterization

The electrochemical behavior of the microelectrodes was studied in PBS (1 mM, pH 7.4) by electrochemical impedance spectroscopy (EIS). For the electrochemical characterization, a potentiostat (Reference 600, Gamry Instruments) was used with a three electrode cell configuration where the probe microelectrodes were connected individually as the working electrode, a platinum wire served as the counter electrode, and an Ag-AgCl (3 M KCl, Gamry Instruments) as the reference electrode. The impedance was measured at frequencies from 1 Hz to 100 kHz by applying a sinusoidal signal with an amplitude of 10 mV.

### In vivo recordings

Before and after each surgery, the impedance magnitude of each electrode was measured using a protocol implemented by the RHD2000 series chip (Intan Technologies) with the probe microelectrodes placed in a dish with sterile PBS (1 mM, pH 7.4) and a reference electrode, Ag-AgCl wire (Science Products GmbH, E-255).

For the surgeries under ketamine, Long Evans rats (400 to 700 g, both sexes) were anesthetized with a mixture of ketamine (60 mg/kg) and medetomidine (0.5 mg/kg), and placed in a stereotaxic frame. At the initial stage of each ketamine surgery, atropine was given to suppress mucus secretion (0.1 mg/kg, atropine methyl nitrate, Sigma-Aldrich). For the surgeries under urethane, rats (400 to 700 g, both sexes) of the Lister Hooded strain were anesthetized with urethane (1.6 g/kg) and placed in a stereotaxic frame. At the initial stage of each urethane surgery, the animal was injected with atropine (0.05 mg/kg), temgesic (20 μg/kg) and rimadyl (5 mg/kg). Ketamine, medetomidine and urethane were administered by intraperitoneal injection, while temgesic and rimadyl were administered by subcutaneous injection. Atropine was administered by intramuscular injection

Anesthetized rodents then underwent a surgical procedure to remove the skin and expose the skull above the targeted brain region. Small craniotomies (2 mm medial-lateral and 2 mm anterior-posterior) were performed above the target area. The reference electrode Ag-AgCl wire (Science Products GmbH, E-255) was inserted at the posterior part of the skin incision. Equipment for monitoring body temperature as well as a live video system for performing probe insertion were integrated into the setup. For the extracellular recordings we used the Open Ephys [27] acquisition board along with the RHD2000 series interface chip that amplifies and digitally multiplexes the signal from the 32 extracellular electrodes (Intan Technologies). Extracellular signals in a frequency band of 0.1-7,500 Hz were sampled at 20 or 30 kHz with 16-bit resolution and were saved in a raw binary format for subsequent offline analysis using the Bonsai framework [28], [29].

Animal experiments under urethane were approved by the local ethical review committee and conducted in accordance with Home Office personal and project (I67952617; 70/8116) licenses under the UK Animals (Scientific Procedures) 1986 Act. Animal experiments under ketamine were approved by the Champalimaud Foundation Bioethics Committee and the Portuguese National Authority for Animal Health, Direcção-Geral de Alimentação e Veterinária.

### Analysis

For the analyses described in the following, a third order Butterworth filter with a band-pass of 250-9,500 or14,250 Hz (95 % of the Nyquist frequency) was used in the forward-backward mode.

The magnitude of the background noise was estimated from the median absolute signal, assuming a normal noise distribution, *σ*_*Median*_ = median(|signal(t)|/0.6745) avoiding contamination by spike waveforms [30]. Alternatively, the noise was defined as the standard deviation (*σ*_*RMS*_) of the signal [13]. Some results are also represented as mean ± standard deviation.

We ran Kilosort [31] on all the datasets with the maximum number of templates set to 128 (four times the number of electrodes on our probe). This algorithm iteratively generates templates and then uses these templates to detect and classify the individual spikes. Each spike is assigned to the template that matches it best. Afterwards, we used Phy [32] to check the automatically generated units/clusters. Phy is a graphical user interface for refining the results of spike sorting. After the manual sorting we used functions to assess cluster quality (https://github.com/cortex-lab/sortingQuality). The “well isolated” units considered for the analysis have simultaneously low interspike interval (ISI) violations and contamination rates, and high isolation distances values. Units with more than 50 spikes were considered for further analyses. Additionally, the average peak-to-peak (P2P) amplitude of all spikes from each unit on a given recording site was computed (see Figure S2 from Supplementary Information).

## RESULTS

### Electrode coating

Figure 1a-c reveals the morphological differences between a pristine iridium electrode, PEDOT-PSS coated electrode, and gold coated electrode (Figure 1a, b, and c, respectively). Pristine electrodes typically display a smooth surface with almost no irregularities (although some might occur during to the microfabrication process). Gold coating creates a rough structure on the electrode, which leads to an increase in surface area, one of the key factors in lowering the impedance modulus at 1 kHz. PEDOT-PSS coated electrodes have a ‘fuzzy’ coating. For gold coated electrodes (Figure 1c and d), we observed that even though the mean impedance after coating is relatively low when compared to the pristine counter-part, these values tended to increase following brain insertion. This may reflect the poor adhesion of the gold coating to the iridium electrodes (Figure 1d). The gold instability and delamination was also observed in some previous studies [13]. In the case of PEDOT-PSS (Figure 1b and e), the impedance values remained stable for a long period of time, allowing for repeated surgeries and penetrations. Therefore, taking into account the impedance value after PEDOT-PSS coating (values under 100 kOhm), the stability and resilience over time, this coating was considered ideal for reducing the polytrode microelectrodes impedance. Figure 2a illustrates the microelectrode array design employed to assess the impact of impedance on data quality (see also Figure S1 from Supplementary Information). Electrodes were coated in a ‘chess board’ pattern such that the signal of one neuron is detected by both coated and non-coated electrodes. The mean impedance at 1 kHz for three polytrodes was 1.1 ± 0.4 MOhm and 0.084 ± 0.015 MOhm pre- and post-coating, respectively. The deposition protocol is stable across probes and electrodes (n_pristine_ = 48 and n_PEDOT_ = 46) (Figure 2b).

**Figure 1.**
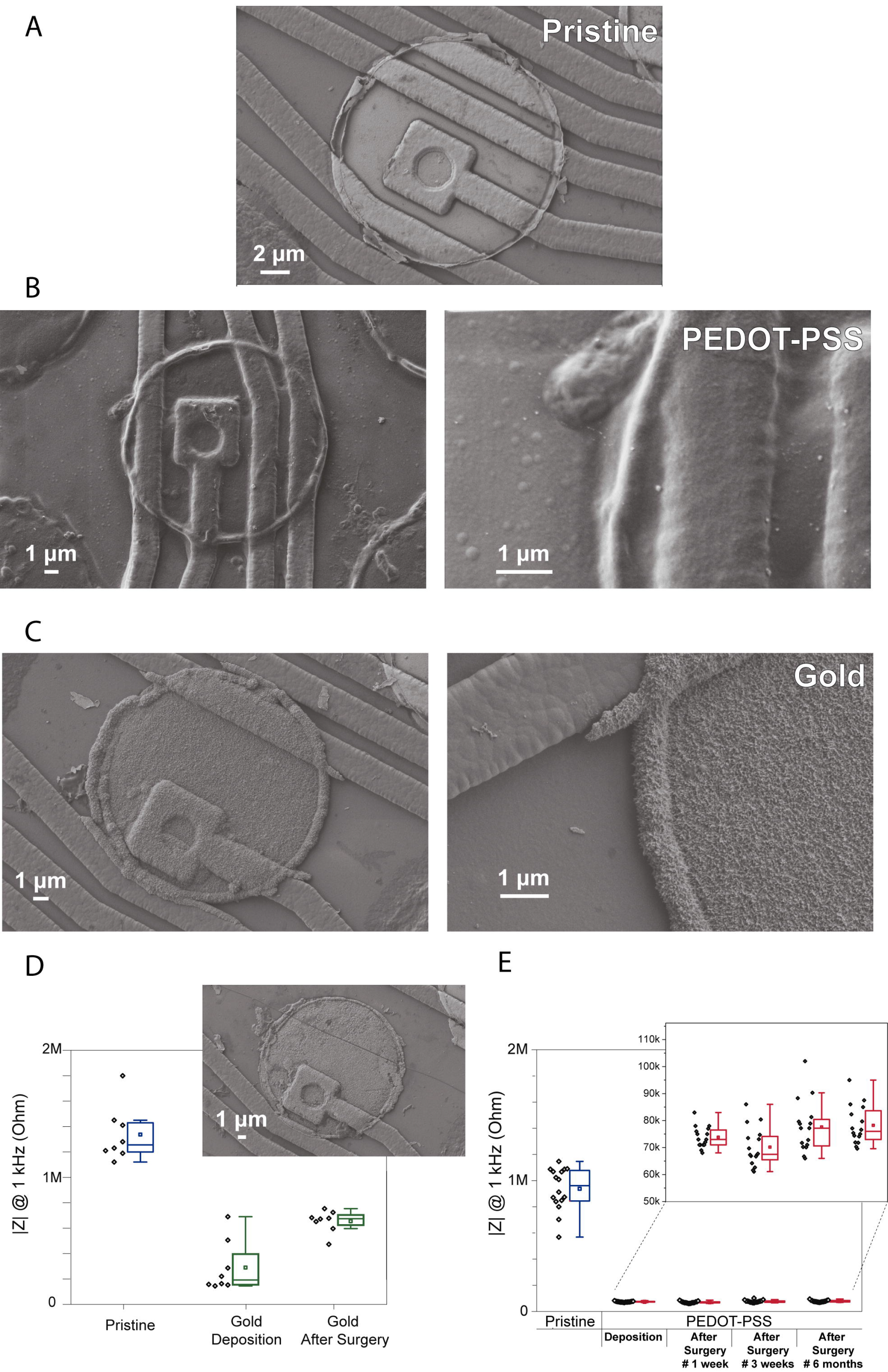
SEM images showing the surface morphology of electrodes from a commercial polytrode in their original state, and after the coatings. (a) Pristine electrode, (b) PEDOT-PSS coated electrode and (c) gold coated electrode; (d) Stability of gold coating for 8 electrodes from one polytrode (impedance variation of electrodes) after the deposition and after an acute surgery. SEM image insert of the gold coating from one electrode after the surgery; (e) Stability of PEDOT-PSS coating for 16 electrodes from one polytrode (impedance variation of electrodes) for the duration of approximately 6 months with surgeries being performed during that period. In the boxplots, line: median, square: mean, box: 1st quartile-3rd quartile, and whiskers: 1.5 x interquartile range above and below the box.

### Noise characterization: in saline

First, the performance of PEDOT-PSS coated electrodes was compared to pristine electrodes in terms of noise, both in saline solution and during *in vivo* recordings. The contribution of all non-biological noise sources was measured by recording signals from single microelectrodes immersed in a saline solution. The non-biological sources include the electronic noise due to the amplifier, thermal noise, and noise associated with the double layer interface [9], [33]. At room temperature, the actual noise measured in saline solution for pristine and coated microelectrodes is shown in Figure 2c. The *σ*_*RMS*_ and *σ*_*Median*_ values are similar in saline solution. The noise in saline, the *σ*_*Median*_ value, for the pristine electrodes was 5.6 ± 0.4 μV, and for the coated electrodes was 3.9 ± 0.4 μV, which represents a reduction of about 30 %.

The thermal noise depends on the real part of the measured impedance and it is defined by 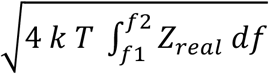, where k is Boltzmann’s constant, T is the temperature in degrees Kelvin, f1 and f2 are the lower and upper limits of the recording bandwidth in Hz, and Z_real_ is the real part of the impedance in the respective frequency bandwidth (f1 to f2). The thermal noise computed in the 200 – 8,000 Hz frequency band for pristine (n = 3) microelectrodes was 5.0 μV and for coated (n = 3) microelectrodes was 2.8 μV (Figure 2d and for a detailed description see Supplementary Information). Additionally, the electronic noise due to the amplifier in our system, measured by shorting the headstage inputs, was 2.0 ± 0.1 μV. We can predict the non-biological noise value as the square root of the sum of the squared thermal noise (~ 5.0 μV and 2.8 μV for pristine and coated microelectrodes, respectively) and squared electronic noise (~ 2.0 μV). We found similar values for the total noise measured in saline (5.6 μV in non-coated and 3.9 μV in coated) and the predicted ones (5.4 μV in non-coated and 3.4 μV in coated).

### Noise characterization: in vivo

We next recorded *in vivo* using polytrodes with the ‘chess board’ pattern described in Figure 2a. These recordings were conducted in different brain regions and at different depths (Figure S3 and Table S1 from supplementary Information). Also, ketamine or urethane anaesthesia was used to compare signal and noise levels recorded during different brain states (Figure 2e and f). Under ketamine, the cortex switches between periods of high neuronal activity and periods of much lower activity (up and down states)[34]. Under urethane anaesthesia, the activity is similar to natural brain activity during sleep [35], [36].

**Figure 2.**
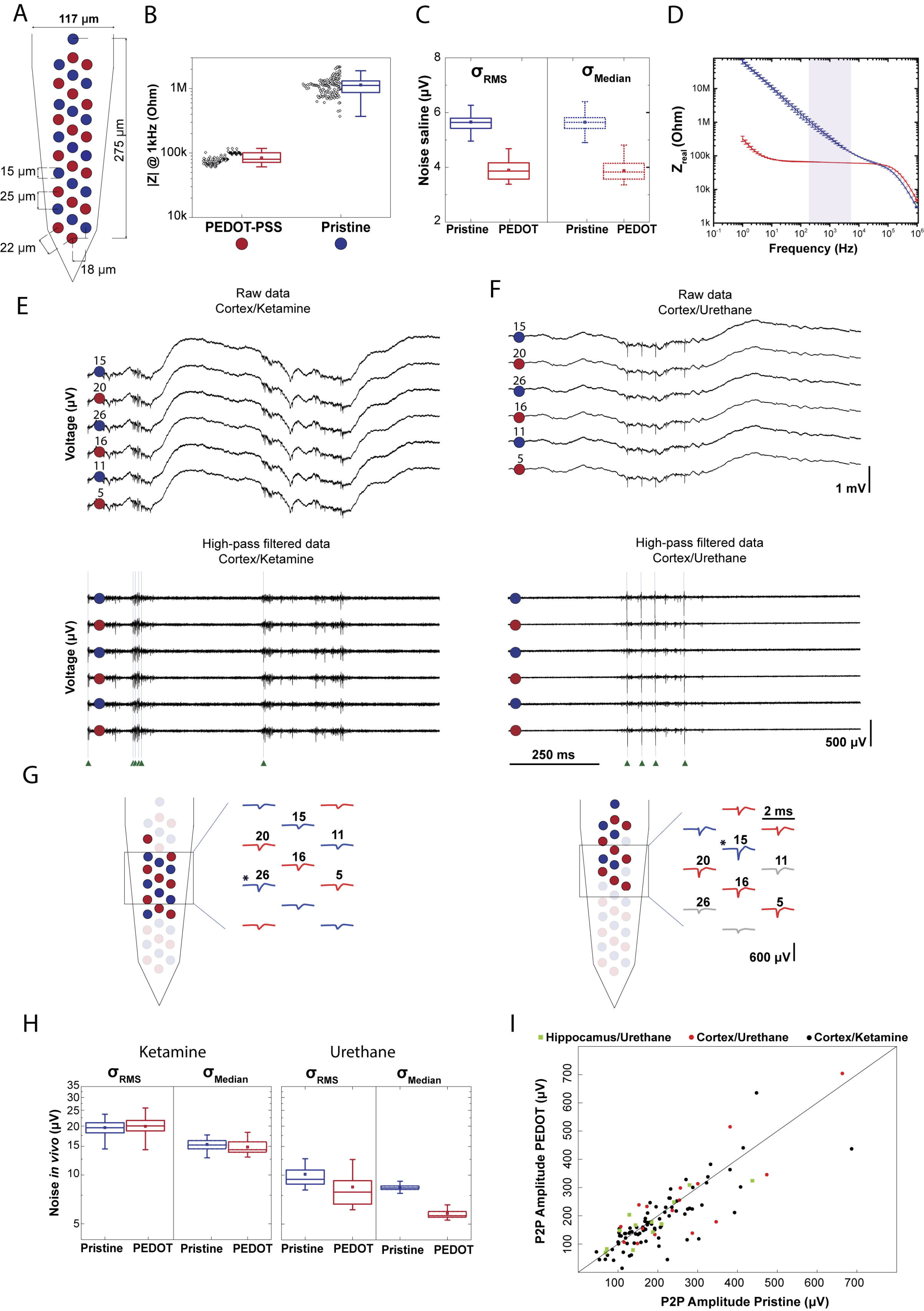
Impact of impedance on data quality. (a) Schematic of a polytrode where electrodes were modified in a ‘chess board’ pattern. Red circles represent PEDOT-PSS electrodes and blue circles represent pristine electrodes; (b) Stability of PEDOT-PSS deposition protocol. Impedance measured for 3 polytrodes (n_pristine_ = 48 and n_PEDOT_ = 46). Black points denote individual measurement for each electrode (3 measurements for each electrode); (c) Noise *σ*_*RMS*_ and *σ*_*Median*_ of recordings performed in PBS (n_pristine_ = 48 and n_PEDOT_ = 46). (d) Impedance spectroscopy of PEDOT-PSS coated (n=3) and pristine (n=3) electrodes shows a significant decrease in the impedance real value. The light purple shaded area corresponds to the frequency range in which the thermal noise was computed; (e)1 s-long raw data traces from 6 electrodes, 3 coated and 3 non-coated, from the recording ‘*amplifier2014_11_25T23_00_08.bin’*. This recording was carried out in cortex under ketamine anaesthesia. Top: signals correspond to the 0.1–7.5 kHz frequency band. Bottom: high-pass filtered traces to highlight spontaneous spiking activity. Green arrows indicate the time of spikes identified for a putative neuron; (f) The same representation as in (e) for the recording ‘*amplifier2017_02_02T15_49_35.bin’*. This recording was carried out in cortex under urethane anaesthesia; (g) Representative putative neurons from each of the recordings shown above. Left panel corresponds to the cortex/ketamine recording and right panel to the cortex/urethane recording. Schematic of two polytrodes with red and blue colored waveforms and circles denoting the electrodes with higher peak-to-peak amplitudes from each unit, respectively. The asterisks indicates the electrode with the highest amplitude P2P; (h) *σ*_*RMS*_ and *σ*_*Median*_ of 9 recordings performed in rat cortex, 6 of wich under ketamine, and 3 under urethane (n_PEDOT_ket_ = 96, n_Pristine_ket_ = 96, n_PEDOT_ure_ = 48 and n_Pristine_ure_ = 48); (i) The maximum P2P amplitude per unit for coated electrodes and for non-coated is plotted. The P2P amplitude averages from 109 clusters are plotted. In the boxplots, line: median, square: mean, box: 1st quartile–3rd quartile, and whiskers: 1.5 x interquartile range above and below the box.

Figure 2e, f and h highlight very different levels of noise *in vivo* due to variations in background neural firing rate (i.e., biological noise level is highly variable). Note that, in general, the levels of noise under ketamine are higher compared to urethane, due to the increase in this background activity. Moreover, the values of noise vary with the method used to compute the noise magnitude. Higher values for the noise *in vivo* were found when taking into consideration *σ*_*RMS*_ values, probably due to a contribution of spikes. The *σ*_*RMS*_ value is based on the standard deviation of the signal, which increases with the firing rate [30]. Therefore, the *σ*_*Median*_ noise values were used to compare the noise between experiments, and within an experiment. Under urethane, the *σ*_*Median*_ values from coated electrodes are smaller compared to the non-coated electrodes. On average, the *σ*_*Median*_ value was reduced from 8.4 ± 0.4 μV in non-coated to 5.8 ± 0.5 μV in PEDOT coated microelectrodes, a ~ 30 % reduction. Under ketamine the *σ*_*Median*_ noise was 15.4 ± 1.2 μV in non-coated and 14.8 ± 1.3 μV in PEDOT coated microelectrodes. The noise values found for *in vivo* recordings are highly variable (Figure 2h) and the noise reduction observed in saline is likely preserved *in vivo*, yet masked by the much larger variation in background spiking activity.

Does the difference in noise observed between coated and non-coated electrodes matter for detecting spikes? Usually, the negative voltage deflection of a well isolated unit exceeds 40-70 μV. Therefore, the benefits resulting from the ~ 2 μV noise reduction achieved by coating electrodes would be largely irrelevant for detecting spikes.

### Signal characterization: amplitude of action potentials

Although not resulting in a major reduction of noise at relevant frequencies, it is still possible that coating electrodes might increase the magnitude of each spike’s signal. Figure 2g shows two examples of putative neurons where each waveform corresponds to the average of all the spikes from each unit on a given recording electrode. Additionally, red and blue colored waveforms and circles denote electrodes where the peak-to-peak average amplitude is larger than half of the maximum peak-to-peak average amplitude of the isolated unit. Therefore, they represent the electrodes with the highest peak-to-peak amplitude from each unit.

For each of the 109 putative neurons sorted from 11 recordings, the largest average peak-to-peak amplitudes from the pristine and PEDOT electrode groups were plotted (Figure 2 i). Therefore, for each unit, two values are plotted in Figure 2i, corresponding to the pristine and PEDOT channel with the largest average peak-to-peak amplitude. If the largest peak-to-peak amplitude spikes are detected by the PEDOT coated electrodes (low impedance electrodes), then the scatter points would fall above the unity line. However, if the largest peak-to-peak amplitude spikes are detected in the pristine electrodes (high impedance electrodes) the scatter points would fall below the line. Our results show that the probability of recording spikes exceeding an amplitude peak-to-peak of 40 μV is similar for coated and non-coated electrodes. Therefore, there is no obvious relationship between impedance and the peak-to-peak amplitude of sorted units in this impedance range (100 kOhm to 1 MOhm).

## DISCUSSION

### Side-by-side impedance comparison

The ability to record from closely-spaced electrodes permitted accurate comparisons between electrodes with two very different impedance values. The PEDOT-PSS deposition protocol made it possible to decrease impedance up to tenfold on average, from 1.1 ± 0.4 MOhm to 0.084 ± 0.015 MOhm. We divided our noise analysis into non-biological noise (noise measured in saline solution) and biological noise, where the level of noise was assessed during acute recordings within the cortex of anesthetized rats. As expected with the impedance reduction, we found a reduction in noise magnitude in saline after coating, since the thermal noise is proportional to the square root of the real part of the impedance [9]. The reduction in impedance resulted in an average ~ 30 % decrease in the non-biological noise. Nevertheless, when using electrodes *in vivo*, this reduction in the thermal noise is largely overwhelmed by the much larger biological noise and would not improve the detection of spikes with commercial polytrodes. Moreover, we found no significant effect of impedance on spike peak-to-peak amplitude and detection probability on both coated and non-coated electrodes. In summary, the impedance values found at 1 kHz in commercial silicon polytrode microelectrodes don’t seem to affect data quality during spike recording. Moreover, the entire dataset used to quantify the effect of an electrode’s impedance on data quality is available online (http://www.kampff-lab.org/polytrode-impedance/) and summarized in Table S1 of the Supplementary Information.

### But why such different views about the role of impedance?

Electrophysiological studies routinely report two different views of the impact of impedance on data quality. Many studies show that decreasing the impedance improves the signal-to-noise ratio, while others find that impedance did not affect the data quality or subsequent analysis. Here we will attempt to rectify these discrepant views.

In studies where researchers use tetrodes and single microwires, lowering the impedance is beneficial because a low-impedance electrode minimizes signal loss through shunt pathways (usually capacitive coupling to ground). Shunt capacitance can be significant in long, thinly-insulated electrode wires that connect a recording site to the pre-amplifier [37]. Thus, for tetrodes and microwires, lowering impedance will result in a larger signal for both local field potentials and spikes [4]. However, with silicon polytrodes, shunt capacitance is much smaller and does not appear to cause signal attenuation for typical values of polytrode electrode impedance [38].

However, if polytrodes, particularly those with higher impedance values (> 2 MOhm), are used with an amplifier that has a (relatively) low input impedance, then a voltage-divider is formed between the electrode and amplifier. The amplifier from Intan Technologies has an input impedance of 13 MOhm, and with electrode impedances of 1 MOhm and 100 kOhm, the signal loss is around 7 % and 1 %, respectively, which may be negligible, but for an electrode with 3 MOhm impedance, this signal loss is around 20 %. For more detailed explanation, see Figure S4 from Supplementary Information.

Do we need to coat our polytrode electrodes? No, assuming we have a good amplifier and low shunt capacitance. But we propose that microelectrode coatings, in chronic applications, may do more than just reduce the impedance. Some coatings may help to promote cell health at the electrode surface and minimize the immune response of surrounding brain tissue. Strong neural attachment to implanted electrodes is desirable as it increases interface stability and improves electrical transfer across the tissue-electrode interface [3], [22], [39], [40]. We thus propose that we stop worrying about impedance magnitude (as long as it stays well below the input impedance of the amplifier) and start focusing on bio-compatible materials [39], [41], [42].

## Supporting Information

Information about the commercial polytrode, an example of one of the 109 putative neurons, a detailed explanation about the thermal noise calculation, information about the acute recordings performed in anesthetized rat brain, summary of the dataset gathered to quantify the effect of electrode impedance, and a detailed explanation about the effect of electrode’s impedance on signal amplitude.

## Author Contributions

J.P.N., G.L., J.F. and A.R.K. conception and design of research; J.P.N., J.F., J.N., and P. Baião performed experiments; J.P.N. and A.R.K. analyzed data; J.P.N., J.F., and A.R.K. interpreted results of experiments; J.P.N. and A.R.K. prepared figures; J.P.N. and A.R.K. drafted manual. All authors have given approval to the final version of the manuscript.

## Note

Readers are alerted to the fact that additional materials related to this manuscript may be found at http://www.kampff-lab.org/polytrode-impedance/.

## ACKNOWLEDGMENT

We would like to thank the institutional support and funding provided by the Champalimaud Foundation, CENIMAT/I3N and Sainsbury Wellcome Centre (funded by the Gatsby Charitable Foundation and the Wellcome Trust).

---

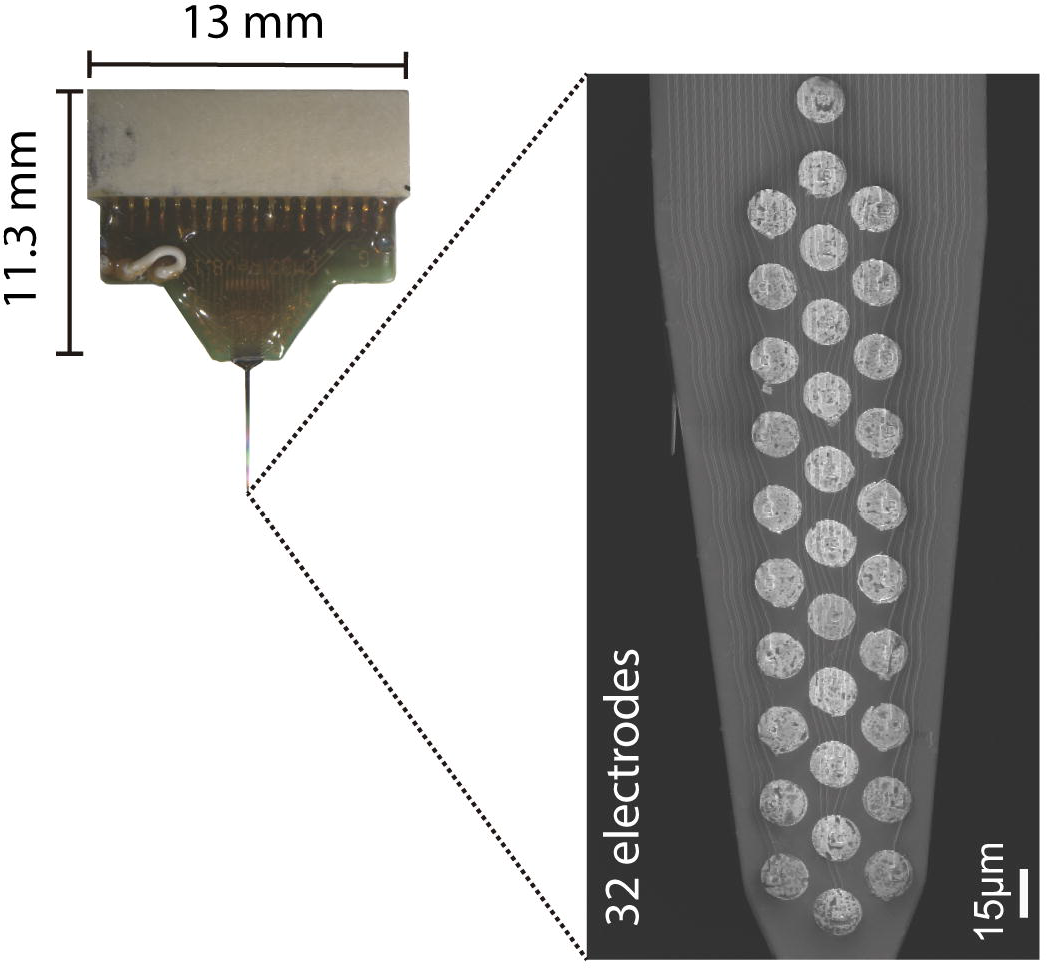

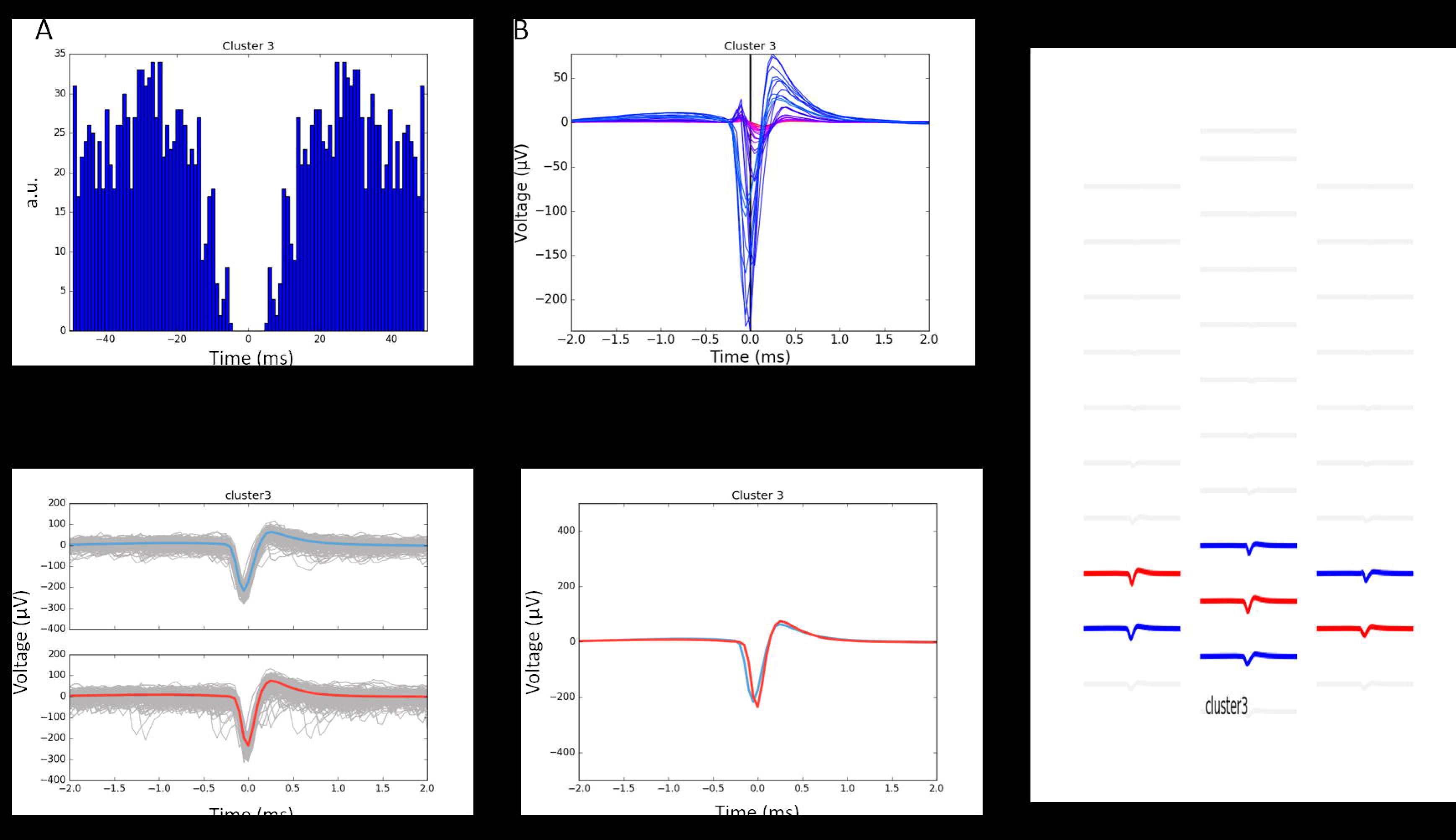

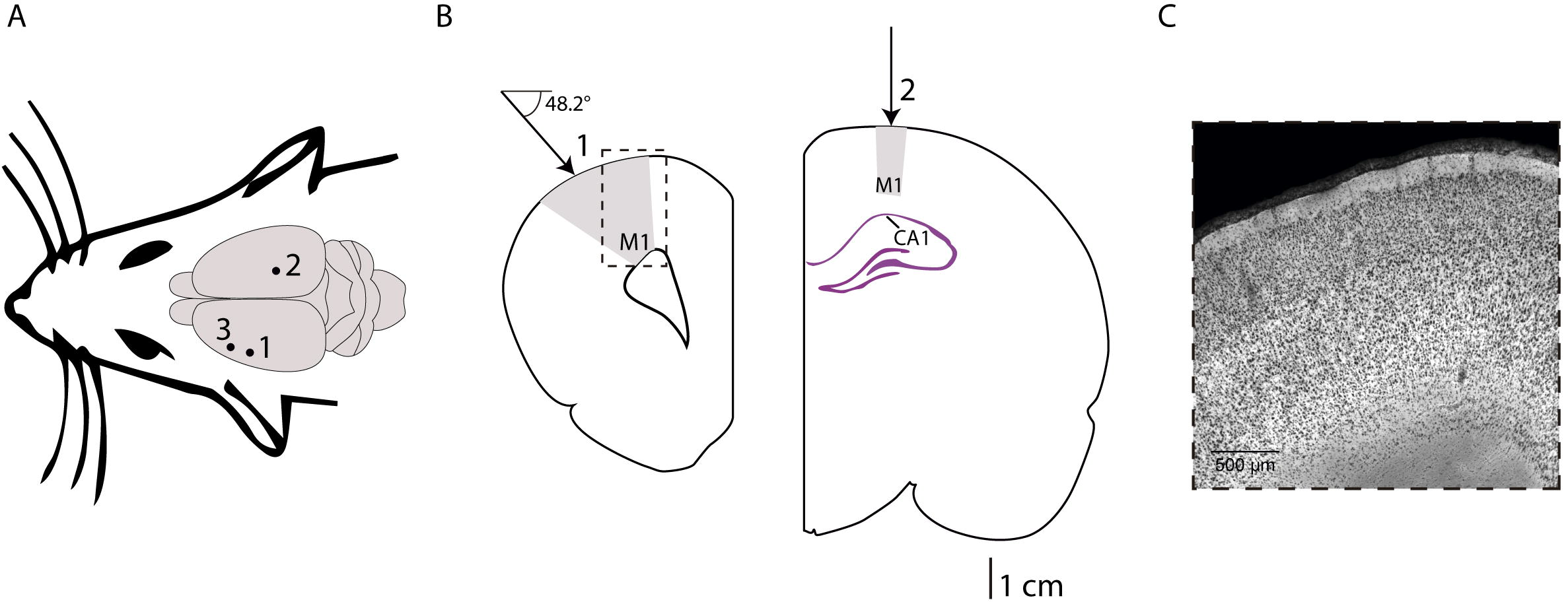

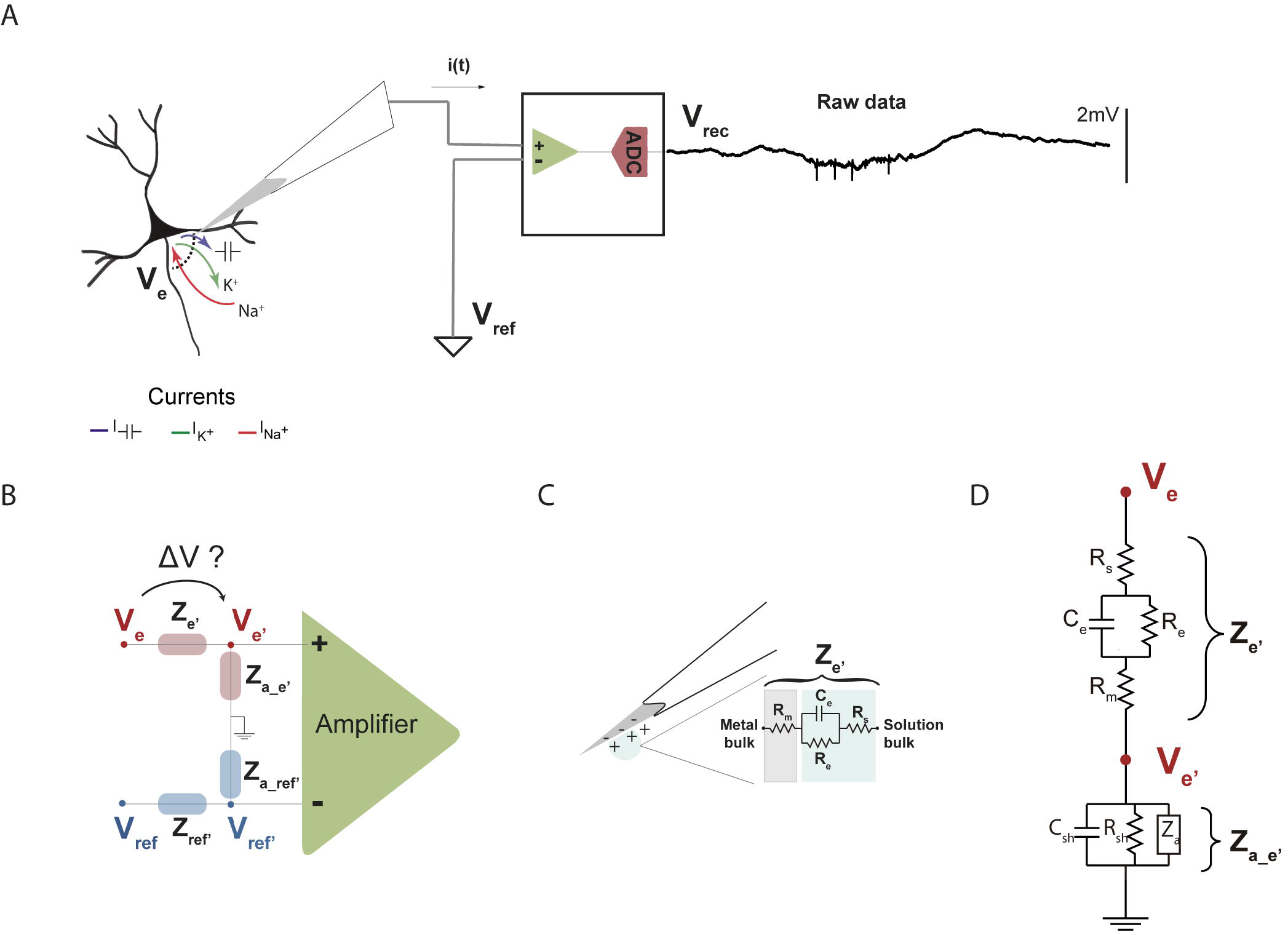

